# A social media-based framework for quantifying temporal changes to wildlife viewing intensity: Case study of sea turtles before and during COVID-19

**DOI:** 10.1101/2022.05.19.492636

**Authors:** Kostas Papafitsoros, Lukáš Adam, Gail Schofield

## Abstract

Documenting how human pressure on wildlife changes over time is important to minimise potential adverse effects through implementing appropriate management and policy actions; however, obtaining objective measures of these changes and their potential impacts is often logistically challenging, particularly in the natural environment. Here, we developed a modular stochastic model that infers the ratio of actual viewing pressure on wildlife in consecutive time periods (years) using social media, as this medium is widespread and easily accessible. Pressure was calculated from the number of times individual animals appeared in social media in pre-defined time windows, accounting for time-dependent variables that influence them (e.g. number of people with access to social media). Formulas for the confidence intervals of viewing pressure ratios were rigorously developed and validated, and corresponding uncertainty was quantified. We applied the developed framework to calculate changes to wildlife viewing pressure on loggerhead sea turtles (*Caretta caretta*) at Zakynthos island (Greece) before and during the COVID-19 pandemic (2019-2021) based on 2646 social media entries. Our model ensured temporal comparability across years of social media data grouped in time window sizes, by correcting for the interannual increase of social media use. Optimal sizes for these windows were delineated, reducing uncertainty while maintaining high time-scale resolution. The optimal time window was around 7-days during the peak tourist season when more data were available in all three years, and >15 days during the low season. In contrast, raw social media data exhibited clear bias when quantifying changes to viewing pressure, with unknown uncertainty. The framework developed here allows widely-available social media data to be used objectively when quantifying temporal changes to wildlife viewing pressure. Its modularity allowed viewing pressure to be quantified for all data combined, or subsets of data (different groups, situations or locations), and could be applied to any site supporting wildlife exposed to tourism.

## Introduction

As awareness of the ecological, economical, and intrinsic value of wildlife has grown in recent decades (König *et al*., 2020), demand to observe animals in their natural environment has also risen (Moorhouse *et al*., 2015). However, this demand (and consequent pressure) is not consistent within or across years (Moorhouse *et al*., 2015), particularly in areas supporting seasonal migrants, such as breeding humpback whales and sea turtles. The recent COVID-19 pandemic exemplifies this issue via the unprecedented absence of ecotourism in hotspots globally, which provided new insights on animal behaviour and movement patterns in the absence of human disturbance (Rutz *et al*., 2020; Bates *et al*., 2021; March *et al*., 2021; Schofield *et al*., 2021). Even within the same population, animals are not randomly distributed in time or space, with certain individuals (in hotspots or residents) being disproportionately targeted (Semeniuk *et al*., 2009; Christiansen and Lusseau, 2014). Thus, while documenting the impacts of wildlife viewing is widespread in both terrestrial and marine environments (Burgin and Hardiman, 2015; Larson *et al*., 2016; König *et al*., 2020), approaches that capture temporal variation in both animal presence and viewing records are required to quantify impacts on individuals, groups and populations (Moorhouse *et al*., 2015; Birk *et al*., 2020; Western *et al*., 2020; Marion *et al*., 2020). By understanding how these pressures change within and across years, relevant actions that promote conservation efforts and mitigate disturbance could be implemented.

The rise of social media, image sharing and the widespread use of mobile phone cameras has generated an extra level of pressure on wildlife, through huge demand for “selfies” and “close-up” images with animals, resulting in close encounters between humans and wild animals that have potential negative impacts (Semeniuk *et al*., 2009; Christiansen and Lusseau, 2014; Lenzi *et al*., 2020; Papafitsoros *et al*., 2021; Molyneaux *et al*., 2021; Van Hamme *et al*., 2021). Yet, because viewing wild animals is intrinsically linked with taking photographs and videos, social media is also being explored as a useful tool to inform conservation science (Dickinson *et al*., 2012; Di Minin *et al*., 2015; Toivonen *et al*., 2019). In particular, wildlife imagery initially uploaded online for purposes other than facilitating conservation studies (e.g. to social media to share personal experiences) is being increasingly applied for science as *passive citizen science* (Edwards *et al*., 2021), *passive crowdsourcing* (Ghermandi and Sinclair, 2019), and *iEcology* (Jarić *et al*., 2020). Such data are being used to quantitatively evaluate interactions between humans and wildlife in both terrestrial (e.g. mountain gorillas (*Gorilla beringei*) (Van Hamme *et al*., 2021), orangutangs (*Pongo abelii*) (Molyneaux *et al*., 2021)) and marine enviroments (e.g. sea turtles (*Caretta caretta*) (Papafitsoros *et al*., 2021), monk seals (*Neomonachus schauinslandi*) (Sullivan *et al*., 2019)).

This application of social media has several advantages over typical researcher-based approaches. For instance, to measure how ecotourism places pressure on wildlife, simple counts of tourist numbers frequenting focal sites (or subset of tourists participating on organised activities) do not necessarily reflect the actual (true) viewing pressure that animals (individuals or groups) are subjected to within a larger population, as operator strategies and animal behaviour and movement patterns also have an effect (Semeniuk *et al*., 2009; Christiansen and Lusseau, 2014). This issue becomes even more challenging when considering nonorganised and nonregulated activities where encounters are often incidental (Papafitsoros *et al*., 2021). In comparison, counting the number of times that animals appear on social media in given time frames could be used to quantify actual viewing pressure on animals more objectively, particularly when exploring how pressure changes over time (seasons/years).

However, systematic approaches to make such comparisons over time based on social media are lacking (Barros *et al*., 2019; Rice and Pan, 2021). Such approaches are needed, because the frequency at which animals are recorded is influenced by the location and number of people accessing social media platforms, plus the availability of cameras and smartphones (Toivonen *et al*., 2019; Ghermandi and Sinclair, 2019). For instance, the number of Instagram (a popular sharing social media platform) users globally has risen from around two million in 2010 to more than one billion in 2020 (https://www.statista.com/statistics/183585/instagram-number-of-global-users/). Consequently, any change in pressure based on comparing the number of animal images uploaded to this platform must be corrected to account for this increase. Challenges also exist because the flow of information from a given human-animal interaction to it appearing in social media is governed by many factors (Tenkanen *et al*., 2017; Jarić *et al*., 2020; Edwards *et al*., 2021). Not all observed animals are captured on camera, nor do all those captured on camera actually appear on social media (Tenkanen *et al*., 2017). Ultimately, the number of human-animal encounters appearing on social media tend to be several orders of magnitude lower than the actual number originally observed (Wood *et al*., 2013; Papafitsoros *et al*., 2021). Even when social media data can be compared temporally (i.e. records are consistent), certainty varies with the number of data samples (observations), with confidence being higher when sightings are higher (consistency of an estimator, Lehmann and Casella, 2006). Thus, it is important to model the uncertainty that exists in this flow of information, and establish the minimum number of social media records required to ensure interpretations have high confidence. Such models are currently absent from the literature for social media, but parallels exist for in other fields (Huang *et al*., 2017).

Here, we developed a rigorous and mathematically consistent framework to quantify temporal changes in the number of human-animal encounters using social media data. We focused on quantifying uncertainty in these changes. We then applied the framework to quantify changes to wildlife viewing pressure on loggerhead sea turtles (Zakynthos Island, Greece) during the course of COVID-19 global travel disruption (2019 to 2021). Because our framework only requires targeted social media mining for photographs and videos of animals in a given focal area, it could be widely applied to any site where humans interact with and photograph wildlife. Through increasing the reliability of using social media-based methods to quantify wildlife tourism pressure, our model facilitates the integration of this global citizen-based phenomenon as a science tool to identify and mitigate adverse effects of human-wildlife interactions objectively.

## Model development and validation

### Real and detected viewing pressures

We developed a rigorous method to compare human viewing pressure on wildlife in a given year against other years, based on detected observations from social media. The following parameters were, first, defined:

- **Real viewing pressure V_real_** number of times that an animal at a focal site is observed by a human. It is difficult to obtain this value because it requires continuous observation of the same animal(s).
- **Photographs of individuals V_photo_:** number of times that a photograph/video of an animal at a focal site is recorded. By definition, this number is larger than or equal to V_real_.
- **Detected viewing pressure V_detected_:** number of times that an animal is recorded for which a person uploaded the image (photograph/video) on a public social media account together with detectable identifiers, e.g. “hashtags” (“#”). We used the term *entry* for such images. By definition this number is larger than or equal to V_photo_.

These parameters were evaluated for all focal animals across all situations, locations and years. However, certain animal groups, observation conditions, locations or years could also be separated to evaluate different types of pressures at different temporal scales. We focused on comparing V_real_ across years using the corresponding V_detected_, which was assumed to be known, following a targeted search in social media.

### Transition probabilities

To link V_detected_ and V_real_, we delineated three *transition probabilities*. When a single observation of an individual animal is made (i.e. one person observes one animal):

P_1_ is the probability that the person took at least one photograph/video of this animal.
P_2_ is the probability that the person has a public social media account.
P_3_ is the probability that given that the tourist has a public social media account, they uploaded a photograph/video of the animal to it with detectable identifiers.

The relationship of V_real_, V_photo_, V_detected_ with P_1_, P_2_, P_3_ is shown in Figure 1. From the definitions above it follows that the approximate value of V_photo_ equals to:

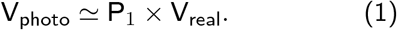

**Figure 1.**
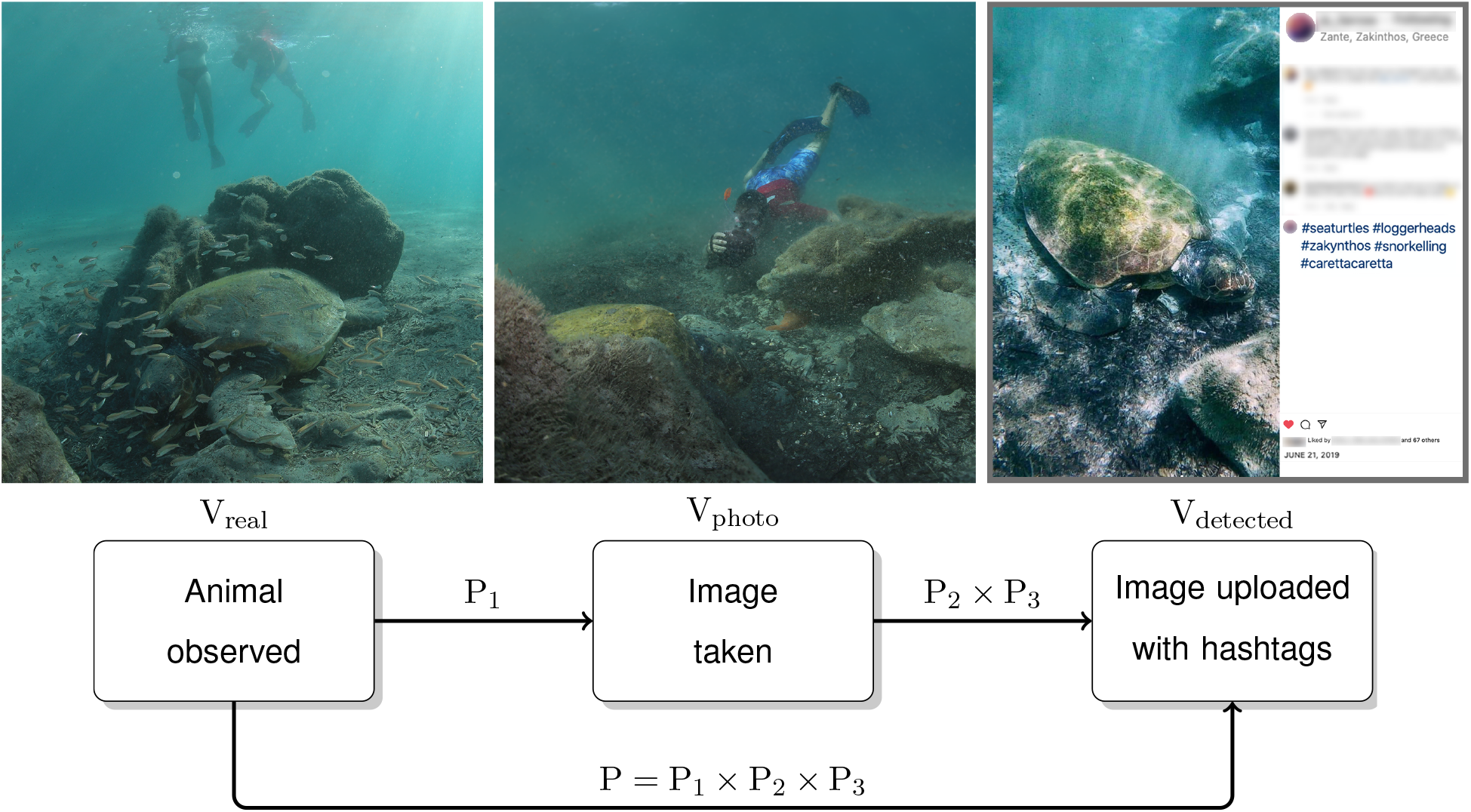
Relationship of V_real_, V_photo_, V_detected_ with P_1_, P_2_, P_3_. Left: actual observation event of an animal, contributing to V_real_. Middle: observer takes a photograph of the animal with probability P_1_, contributing to V_photo_. Right: Given this, the photograph appears on a public social media account with detectable hashtags and probability P_2_ × P_3_, contributing to V_detected_.

Thus, P_1_ is used to transfer from V_real_ to V_photo_. To transfer from V_photo_ to V_detected_, we separated the transition probability into whether the photographer (1) has a public social media account and (2) uploaded the photograph/video with detectable identifiers. The probability that both components are satisfied is equal to P_2_ × P_3_. Thus, P_2_ × P_3_ is used to pass from V_photo_ to V_detected_, meaning that V_detected_ is approximately equal to:

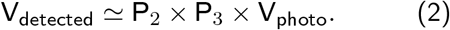

By combining equations (1) and (2) we end up with V_detected_ being approximately equal to

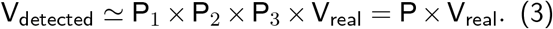

where P denotes the product of the three probabilities. Some assumptions must be made on the transition probabilities:

#### ID-independence assumption

We assumed that P_1_, P_2_, P_3_ do not depend on the identity of the observed animal. For instance, certain individuals are not expected to have higher or lower probabilities of being photographed/videoed than other individuals or for these images to be uploaded on social media.

#### Time-independence assumption

We assumed P_1_, P_3_ would be constant across time. For instance, a person is as likely to record the viewed animal on a device with a camera in all years. The assumption of time independence for P_1_ implies that the availability of photographic equipment (mobile phones, cameras, underwater cameras) remains (approximately) stable across the selected timeframe.

In contrast, P_2_ is considered time dependent because social media users increase annually, and so the probability that a person has a social media account increases, thus we used the notation 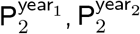 etc.

#### Condition of observation-independence assumption

We assumed that P_2_ and P_3_ do not depend on the *condition* in which the observation took place. For instance, a person would upload photographs/videos with the same probability, regardless of the condition (where or how) in which the observation took place at the focal site. Different conditions could be locations (underwater vs boat based for marine life), means of recording (normal cameras vs drones), or time of the day (daytime vs nighttime). However, in certain situations (i.e. when viewing marine wildlife), P_1_ is likely condition-dependent, with more images being obtained from boat viewings compared to underwater viewings, as devices with cameras that can be used underwater tend to be more expensive and so less used. Thus, we used the notation 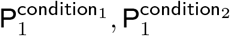 etc.

#### Outcome of viewings-independence assumption

We assumed that the viewing of an animal resulting in a photograph/video that is uploaded to social media does not depend on the corresponding event of a different viewing. However, for this assumption to be true, a person would have to observe just one animal, which is not necessarily the case. This assumption can be validated by quantifying the number of entries uploaded to separate social media accounts. At sites where this is not the case, the model must be interpreted with caution.

### Linking detected viewing pressure to real viewing pressure and related uncertainty quantification

The previous section argued that V_detected_ equals approximately P × V_real_, where P := P_1_ × P_2_ × P_3_. To be more precise from a modelling perspective, V_real_ is a deterministic quantity, whereas V_detected_ is a *random* variable that follows the *binomial* distribution with parameters V_real_ and P, denoted as Bi(V_real_, P). This means that V_detected_ is the number of *successes* (with *success* defined as the event in which a photograph/video was taken and uploaded with hashtags) of V_real_ independent *experiments* (with *experiments* defined as animal viewings), each of which has success probability P. We stated that V_detected_ equals *approximately* to P × V_real_ because the *expected value* of V_detected_ equals to latter; however because V_detected_ is distributed around its expected value (with nonzero variance), its true value is not known. We can only observe its realisation, denoted by v_detected_, which is the number of detected images in social media. This distinction is important from a modelling point of view. The conceptual difference between V_detected_ and v_detected_ can be understood, if a hypothetical experiment is “repeated” by “going back in time”. Even though the probability that V_detected_ belongs to a certain interval remains the same, its realisation v_detected_ would differ due to randomness.

Given some detected entries in two years, 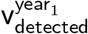 and 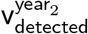, referring to a given time window, condition of observation and group of animals, we wanted to quantify the uncertainty of the corresponding real viewing pressure change, i.e. how much larger/smaller is 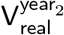 compared to 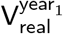. Given an acceptable level of error 0 < *α* < 1, we aimed to find an “interval *I*”, such that 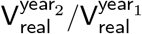 belongs to the interval *I* with probability 1 – *α*. Specifically, we showed that the confidence interval *I* is defined by the following inequalities

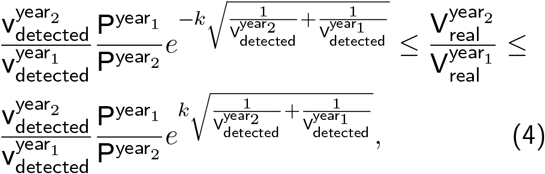

where, *k* is the value of the quantile function of the standard normal distribution at level 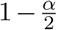. For instance for *α* = 0.05, which we fix henceforth, we have *k* = 1.96. Supplementary material A shows how the confidence interval (4) was derived, and associated justification. In brief, the confidence interval was derived using three assumptions: (i) relatively large values for 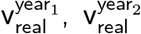, (ii) relatively small values for 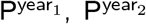 and (iii) the observed values 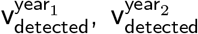 being close to their respective means. Supplementary material B explains these assumptions.

The confidence interval (4) implies the point estimate (most probable value) for real pressure ratio 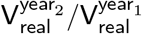 is equal to

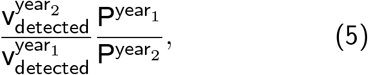

which is natural, as it is simply the ratio of the detected pressure 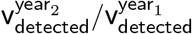 adjusted by the ratio of probabilities 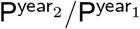. Even though the true probabilities 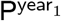 and 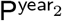 are not known, due to our assumptions, we estimate their ratio as

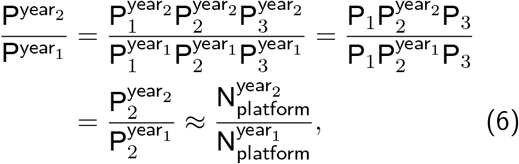

that is, as the ratio of the global number of users of the fixed social media platform in consecutive time frames, i.e. number of users in year_1_ and year_2_, denoted by 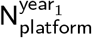 and 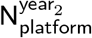 respectively.

### A simple illustration of the confidence interval

To illustrate the confidence interval (4), we provide an example with 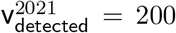 and 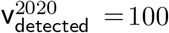 as the number of detected entries. If we assume that the number of social media users did not change, (i.e. 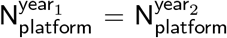), then the confidence interval based on formula (4) is calculated as

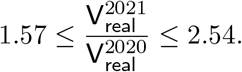

If the number of observations increases (e.g. 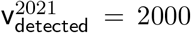 and 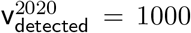) then the new confidence interval shrinks to

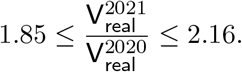

In both cases, the confidence interval lies around the point estimate 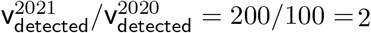 due to (5). As the number of entries increases, the confidence interval shrinks, increasing its robustness and confidence we have in it. However, if the number of social media users doubled, then the confidence interval would shrink by a factor of two to

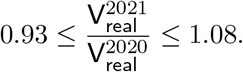

This is logical as the same real pressure with two times the number of Instagram users should result in twice as many entries and consequently double the observed pressure.

### Confidence interval validation

Because three additional assumptions were required to derive the confidence interval (4) we verified its accuracy and validated its use. We computed the confidence interval (4) for a series of 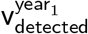 and 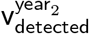 and compared the outputs with the true confidence interval, computed via simulations. Supplementary material D presents the procedure used to compute the true confidence interval. In brief, it was based on the knowledge of transition probabilities, which are not known for real data, and were chosen as 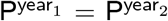 for the simulation. Once we computed these two confidence intervals, we computed their IoU (intersection over union) as a comparison metric. IoU=0 represented no intersection (our estimated confidence interval is not accurate), and IoU =1 represented a perfect alignment (our estimated confidence interval is perfectly accurate).

Figure 2 presents the outputs of this procedure based on 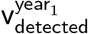 and 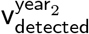 for all possible combinations of numbers between 1 and 50. If either number of detected entries 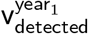 and 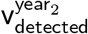 is small, then our estimated and true confidence interval had small IoU, and the estimated uncertainty is not trustworthy. However, this discrepancy is rectified when the number of detected entries rises. Figure 2 (right) quantifies this and shows the minimum number of detected entries 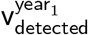 and 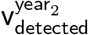 so that IoU reaches a certain threshold. For example, 5 entries are sufficient to obtain an IoU of 75%, whereas 10 entries produce an IoU of 86%. Thus, the results are trustworthy once the number of entries for both are greater than or equal to 5.

**Figure 2.**
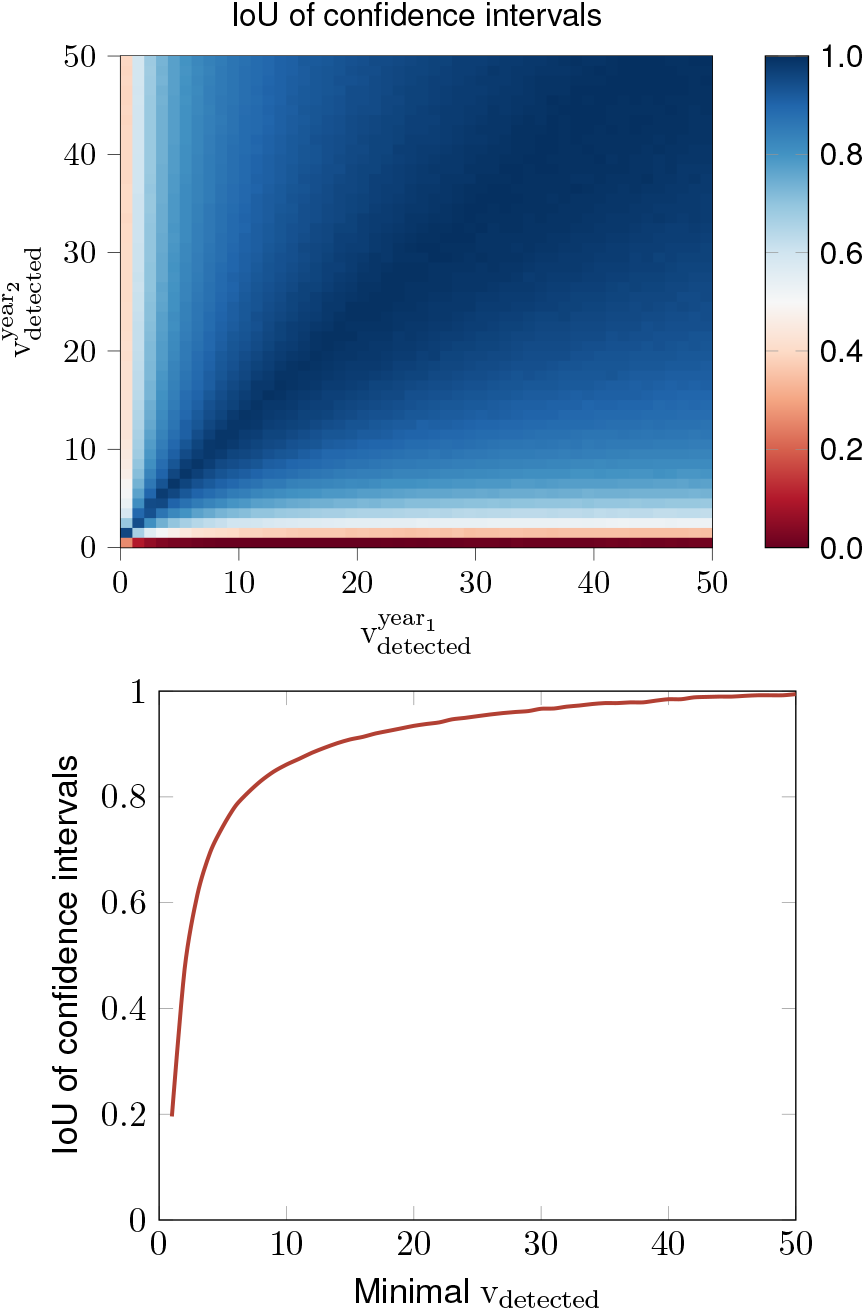
Left: Comparison of our estimated confidence interval (4) and the true confidence interval computed via simulations for 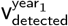 and 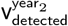 between 1 and 50, using IoU (intersection over union as the similarity metric). Right: IoU of estimated and true confidence intervals as a function of the minimum number of detected entries, with Minimal 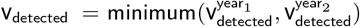.

## Empirical application of the model

### Study site

We tested the model by applying it to a site (Laganas Bay, Zakynthos Island, Greece) supporting large numbers of loggerhead sea turtles (*Caretta caretta*) subject to intense ecotourism (Figure 3). The site supports around 300 breeding adult females seasonally (April-August) (Margaritoulis, 2005; Margaritoulis *et al*., 2011; Schofield *et al*., 2017) and around 40 year-round residents of mostly juveniles and adult males (Schofield *et al*., 2020; Papafitsoros *et al*., 2021). The island is a popular holiday destination in summer (May-October) with over 850,000 visitors. There is a well established wildlife-watching industry, where tourists observe turtles on organised boat tours (Figure 3). This industry is estimated to service 180,000 tourists over 9,000 trips per year, generating an annual revenue of more than 2.7 million euros (Schofield *et al*., 2015; Papafitsoros *et al*., 2021).

**Figure 3.**
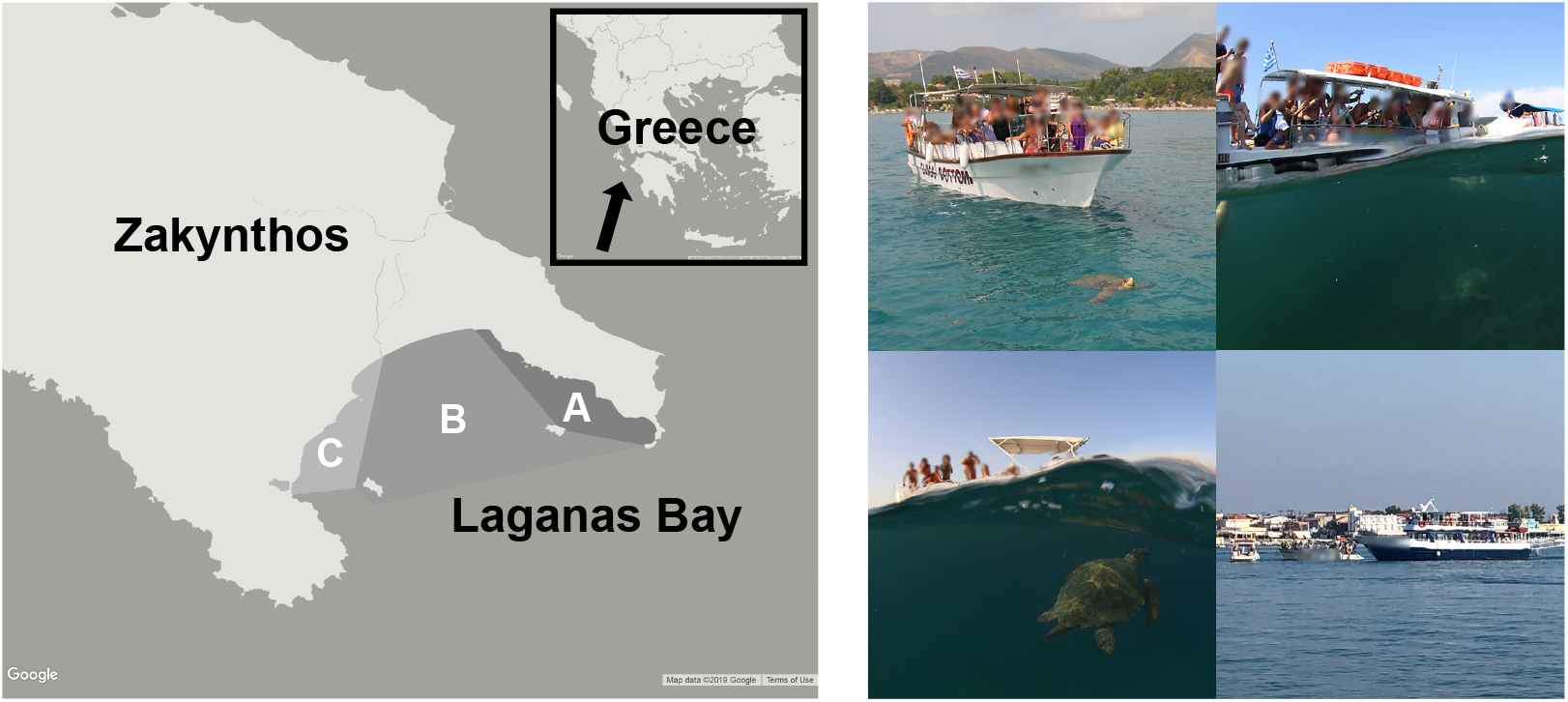
Left: Southern part of Zakynthos Island (Greece) showing Laganas Bay and the maritime zoning of the National Marine Park of Zakynthos (Zones A-B-C). Right: Wildlife watching vessels observing sea turtles. Photograph credits: Kostas Papafitsoros.

### Collection of social media records

Social media records were collected for 2019, 2020 and 2021 following the methods described in Papafitsoros *et al*. (2021). The first year represented a typical tourism year (2019) and the next two years (2020 and 2021) were heavily and mildly impacted by COVID-19, respectively, when tourism levels were limited by global travel restrictions (Schofield *et al*., 2021). Instagram was selected over other social media types because of its popularity and convenient search framework via “hashtags” and “places”. We searched for entries using specific hashtags and places related to the study site and species. The list of hashtags used is provided in the supplementary material of Papafitsoros *et al*. (2021). For an entry to be detected, the user must have a public profile on Instagram and the entry must be uploaded using one of the specified hashtags. The search was performed on at least once weekly from 1 May to 31 October each year, with regular retrospective searches to minimise missed entries. We did not record entries from previous seasons or those used as advertisements by tour operators. For each entry we recorded the date that it was uploaded. We previously showed that more than 80% of entries are uploaded within 2 days of being captured (Papafitsoros *et al*., 2021). We only selected entries taken from a boat, i.e. organised tours, and private hire boats, which defines the “condition of the observation”.

### Number of Instagram users and visitors to Zakynthos

The number of annual Instagram users was obtained from the Statista website^1^, see also Supplementary table 3. The number of arrivals to Zakynthos airport were obtained for each month (May-October 2019-2021) from the Hellenic Civil Aviation Authority (CAA, http://www.ypa.gr). For 2019-2020, the number of daily airport arrivals was also available. Because daily airport arrivals in 2021 were not available, we employed a mass conserving interpolation approach to generate a series of simulated daily arrivals, the sum of which (over all days) is the same with the original sum (over months)^2^. This approach was validated using 2019 and 2020 airport arrivals for which the real number of daily arrivals was available.

### Comparing real viewing pressure ratios with baseline values

We investigated how our developed framework provides more reliable information on changes to viewing pressure from year_1_ to another year, (i.e. year_2_). We compared the estimated ratio of V_real_ with the corresponding confidence intervals to two baseline values (constant 1 and ratio of visitor arrivals). Constant 1 represented equal viewing pressure in year_1_ and year_2_. When the ratio of V_real_ is above (below) value 1, then viewing pressure increased (decreased) from year_1_ to year_2_. The second baseline value is the ratio of visitor arrivals. We consider the ratio of the airport arrivals to be representative of the ratio of the total number of people present at this focal site and observing sea turtles. Comparison of the V_real_ ratio to arrivals tests to what degree V_real_ is directly proportional to the number of visitors. When the ratio of V_real_ is not equal to the arrival ratio, changes to V_real_ might also be driven by additional factors, not just the change in the number of tourist arrivals (e.g. more organised tours operating between years). When the ratio of V_real_ is above (below) the arrival ratio, this means that changes to V_real_ are higher (lower) than that predicted by changes to tourist numbers when assuming a simple linear relationship between the number of visitors and number of animal viewings.

### Overview of Instagram records and tourist numbers

We recorded 2646 Instagram entries from boats for 2019, 2020 and 2021 (n=1382, n=387 and n=877 respectively; Figure 4; Supplementary table 1). The total number of visitors in 2019 was 1,288,651 (airport and port combined) versus 386,756 (70% lower) in 2020, due to COVID-19 pandemic travel restrictions. In 2021, there were 803,868 visitors (107.8% higher than 2020, but still 37.6% lower than 2019; Supplementary table 2).

**Figure 4.**
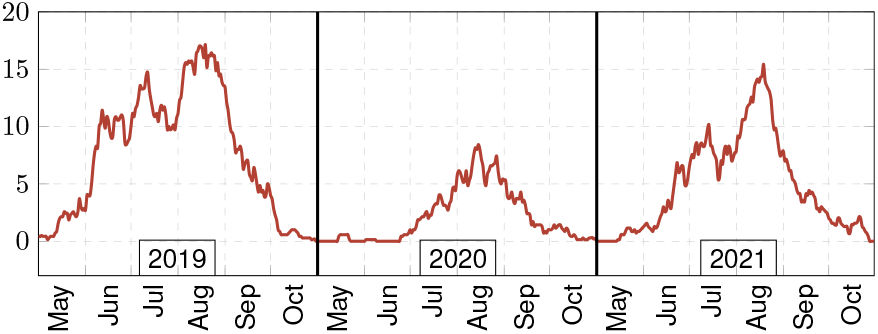
Number of recorded daily Instagram entries taken from boats for May-October for 2019-2021 (moving mean of 7 days).

### Model versus raw data

We investigated three quantities as estimates for the ratio of the ratio 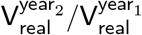:

- raw data ratio 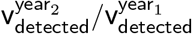;
- point estimate (5) ratio 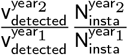;
- associated confidence interval (4).

The raw data ratio was a natural candidate for estimating the real pressure ratio. However, unlike the other two estimates, changes to the number of Instagram users and uncertainty of that estimate were not considered. Therefore, confidence interval (4) might be more reliable than the raw data ratio estimate.

For example, we compared boat viewing pressure before (2019) and during the COVID-19 pandemic (2021) for these three quantities when aggregating viewing images at 3 and 7 day intervals (Figure 5). For the 3 day aggregation window, the raw data ratios 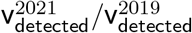 were primarily below 1, apart from three time windows (mid-May, late June, mid-July), suggesting that boat viewing pressure rose from 2019 to 2021 for these specific 3-day windows. However, when adjusting for changes in the number of Instagram users, the corresponding point estimate value of these 3-day windows decreased close to 1, with a high confidence interval above and below 1. Thus, the number of data (detected entries) was insufficient to make objective inferences for this narrow time interval. In contrast, when aggregating viewing images to 7 day intervals, the confidence intervals for June-September remained below 1, indicating that boat pressure change could be inferred with high certainty, and that it was lower in 2020 than 2019.

**Figure 5.**
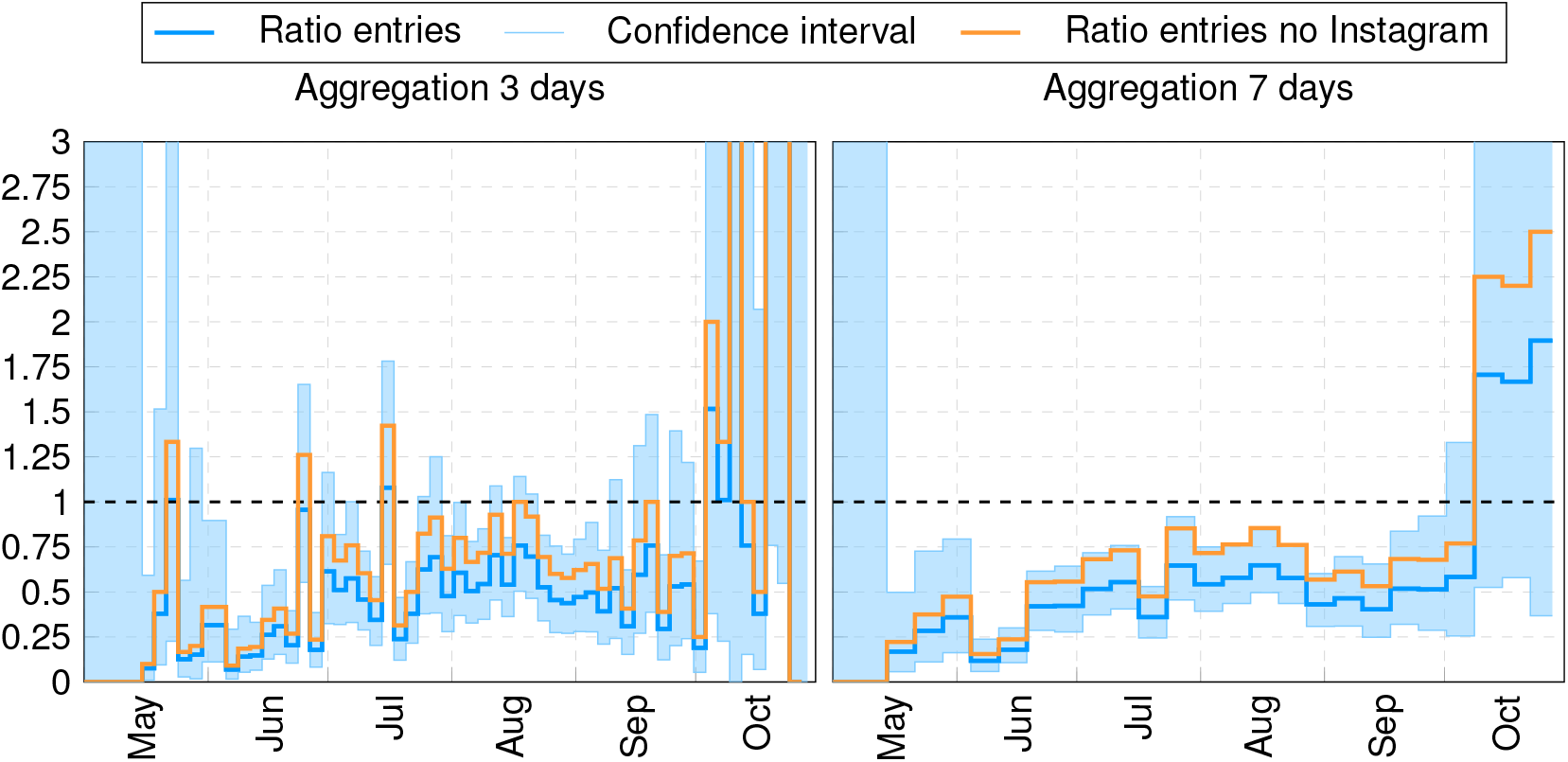
Confidence intervals (light blue) for 2021/2019 ratios of the real boat pressure 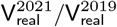 for loggerhead turtles (aggregating viewings in 3 and 7-days intervals). Dark blue line represents the best point estimate, 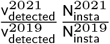; orange line represents the simple ratio of detected entries 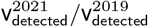 i.e., not accounting for changes in the number of Instagram users. Horizontal dashed line represents equal real viewing pressures for the two years (ratio equal to 1). See https://github.com/sadda/Turtles_Covid for interactive plot for all year ratios 2020/2019, 2021/2019, 2021/2020.

### Effect of time aggregations on the uncertainty of real pressure ratio

To evaluate how aggregating image records in different time intervals impacted uncertainty in the real pressure ratio, we evaluated different time windows (1, 7, 15, 184 days (entire season); Figure 6) for 2020/2019 and 2021/2019. Data based on daily records led to extremely high uncertainty for the estimated real pressure ratio (very large confidence intervals), and should not be used (Figure 6). Certainty increased as the data were aggregated into larger time intervals, with 7 and 15-day intervals providing higher confidence for July-September, since the corresponding detected entries were higher (Figure 7), with high certainty (real pressure ratio below 1), as all confidence intervals were below that number. The arrival ratio closely followed the real pressure ratio, indicating proportionality between real boat pressure and the number of tourists frequenting the site. Based on the entire season (184 days), compared to 2019, boat observation pressure was 25% and 50% lower in 2020 and 2021, respectively, with very high certainty, due to the small confidence interval.

**Figure 6.**
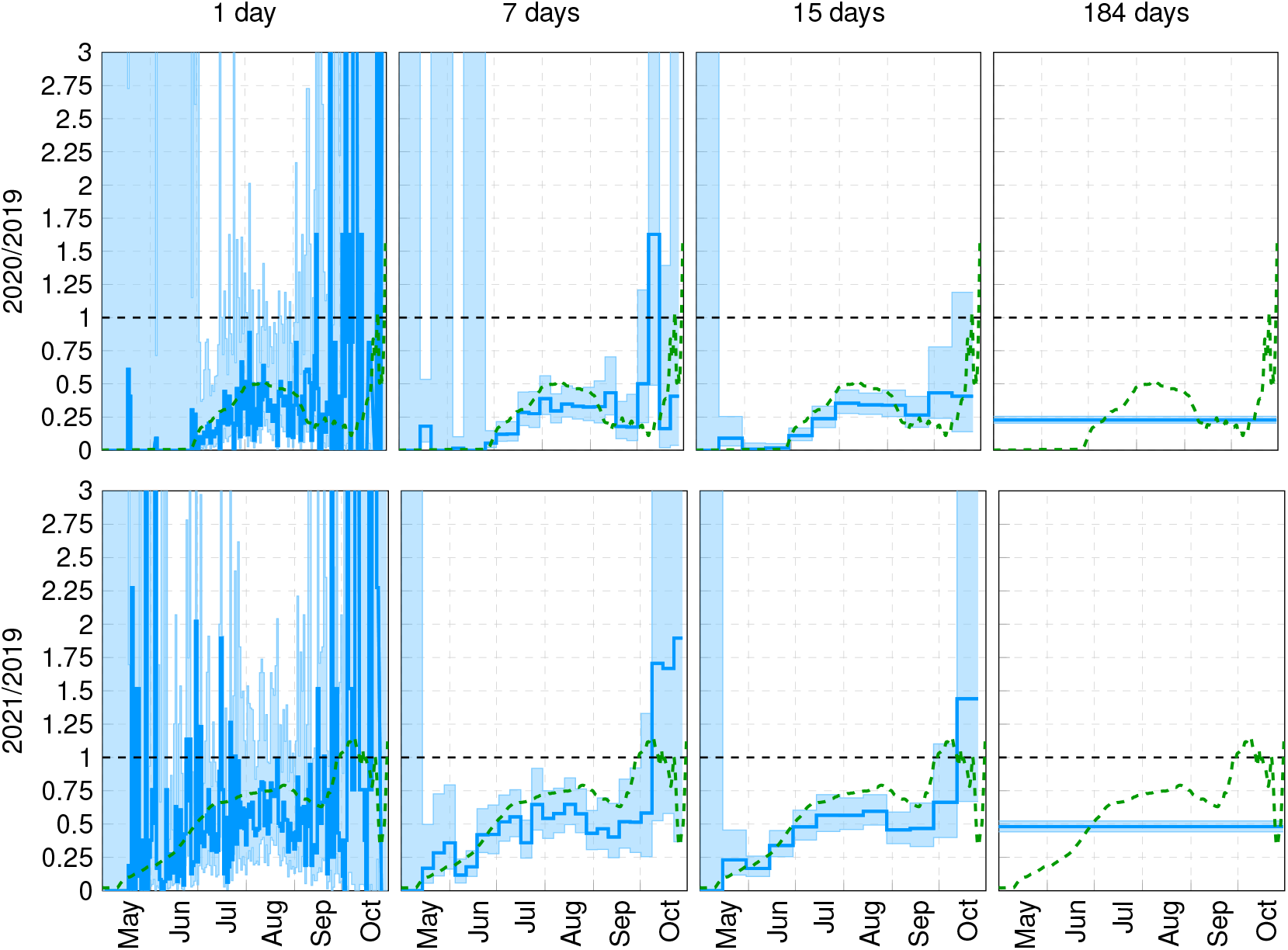
Confidence intervals (light blue) for 2020/2019 (top) and 2021/2019 (bottom) ratios of real boat pressure on loggerhead turtles, 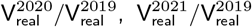 for a series of 1, 7, 15 and 184-day (full period of May-October) aggregation windows. Dark blue line represents the best point estimate. Green line represents the corresponding ratio of visitor arrivals (moving mean of 7 days). Horizontal dashed line represents equal real viewing pressure for the two years (ratio equal to 1). See https://github.com/sadda/Turtles_Covid for interactive plot of all year ratios 2020/2019, 2021/2019, 2021/2020.

**Figure 7.**
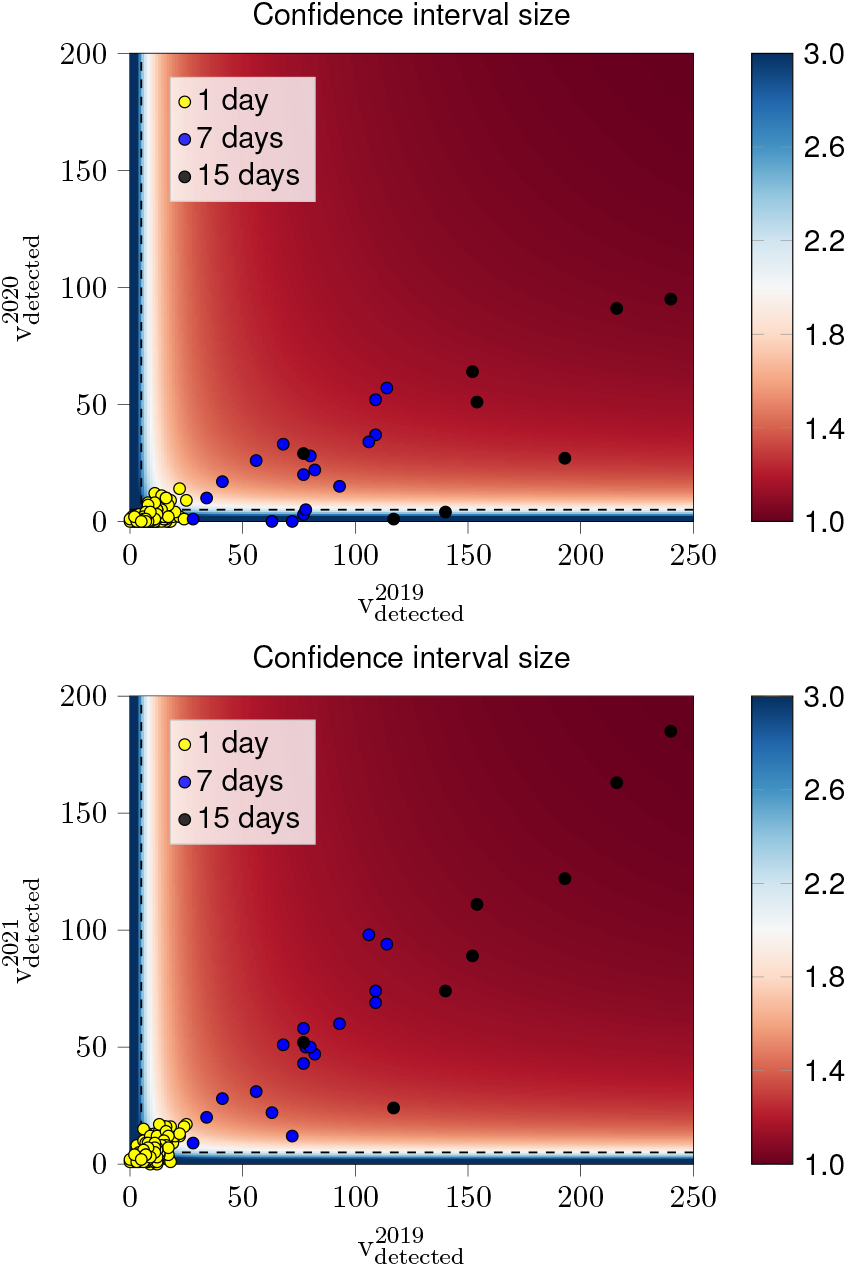
Scatter plot of the number of detected entries for 1-, 7- and 15-day intervals corresponding to the plots of Figure 6.

Figure 8 provides a further visualisation on how the size of the confidence intervals varied with the aggregation window (number of days). We aggregated the days of each month based on this window. For example, for a 4-day aggregation window, a month with 30 days was split into 7 windows (excluding the last 2 days). Then, for each month (May-October), we computed the average size of the confidence interval for each window, defined as the ratio upper over the lower bound of the interval, with 1 corresponding to the smallest possible confidence interval (Supplementary material C). During July-September (and June 2021/2019), the confidence interval had a smaller average width than October, due to the larger number of detected entries. In parallel, the width decreases in all months as the aggregation window increases. For July-September (high visitor season) of 2021/2019, a 7-day aggregation window represents an optimal balance between achieving high certainty (confidence interval size <1.5) with sufficiently fine time scale resolution, since aggregation windows of >7-days do not cause the width of the confidence interval to significantly decrease further. In contrast, a 15-day window was needed for July-September for 2020/2019 ratio (and June 2021/2019), due to fewer entries. For October 2020/2019 and 2021/2019, the same certainty was achieved by setting an aggregation window of >30 days.

**Figure 8.**
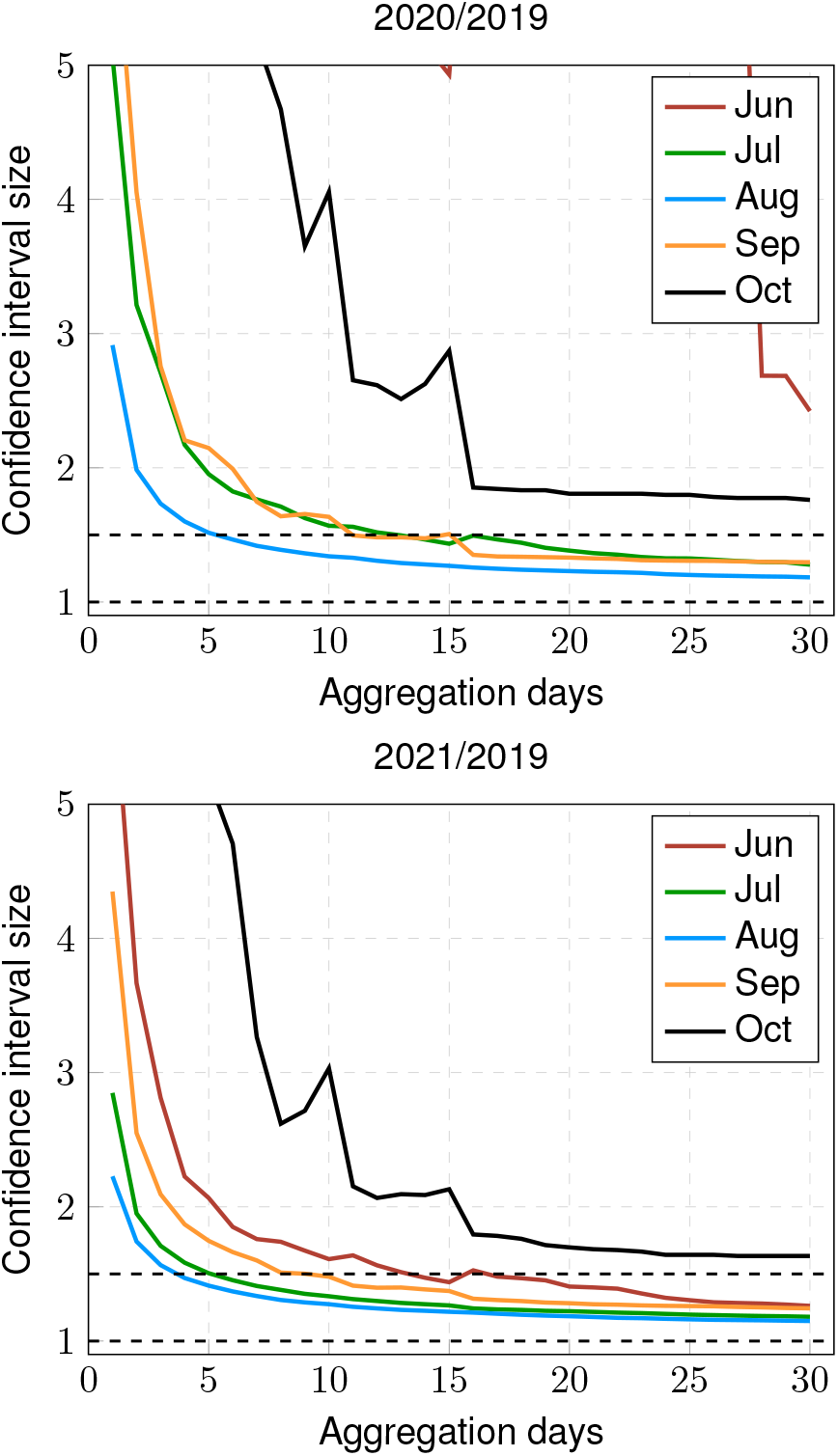
Average confidence interval size (defined as upper bound/lower bound ratio) for real boat pressure ratio in 2020/2019 and 2021/2019 versus different day aggregation windows. Aggregation windows were plotted separately for each month (June-October). May was excluded due to insufficient data.

## Discussion

A key issue of passive citizen science/iEcology raw data and associated analysis is determining uncertainty to improve objective interpretation (Isaac *et al*., 2014; Jarić *et al*., 2020). As such, the present study provided a rigorous framework for using social media imagery with confidence to infer temporal changes (within and across years) in wildlife watching pressure. We showed that confidence increased when integrating multiple days. This framework modelled the flow of information from a human-animal interaction event to that event appearing in social media in a detectable way. This was achieved by introducing the notions of real and detected viewing pressure, linking them via transition probabilities and showing how temporal changes in the former can be estimated by temporal changes in the latter. Detected viewing pressure was modelled as a random variable, allowing uncertainty in real viewing pressure change to be quantified by rigorously deriving confidence intervals. Through applying the model at a site supporting large scale sea turtle ecotourism, we demonstrated its advantages over simply using raw data (i.e., simple ratio of detected animal entries in given time periods). Through increasing the reliability of using social media-based methods to quantify wildlife tourism pressure, our model facilitates the use of social media as a scientific tool providing evidence based information on human pressure.

Our study delineated appropriate time-windows to analyse social media data temporally, allowing robust comparison across years, despite highly different visitor levels due to COVID-19. The optimal aggregation window (balance of high confidence and high temporal resolution) for images at our study site was 7 days during the peak period, with more days being needed during the low periods, due to fewer images. Our results supported those of Tenkanen *et al*. (2017), who showed that social media captured temporal variation in national park visitation rates at a monthly scale, but was challenging on a daily basis, due to fewer social media records. Other studies arbitrarily grouped information into monthly periods. For instance, Molyneaux *et al*. (2021) compared the monthly number of photographs posted on Instagram before and after the onset of the COVID-19 pandemic to quantify variation in interactions between tourists and orangutans. Barros *et al*. (2019) observed that Flickr geotagged data have enough information to capture daily, weekly and monthly distribution patterns of visitors of Spanish national parks, but did not perform temporal comparisons of visitor numbers across years. By implementing our approach, studies could maximise social media data by selecting appropriate aggregation windows of sufficient temporal granularity, guaranteeing certainty in interpretations.

This modularity of our model allows more refined versions to be developed as knowledge of its key constituents becomes available. For instance, estimates of the annual ratio of social media users could be improved by directly incorporating the number of users that belong to the main demographic characteristics (nationality, age group, culture) of visitors to a focal site (Väisänen *et al*., 2021). The time-independence assumption of transition probability P_1_ could also be refined (i.e. availability of photographic equipment across years) by incorporating information on the yearly ratios of global (or more refined demographically) sales of equipment (e.g. smartphones/underwater cameras associated to marine wildlife viewing). Our model could easily incorporate such refinements by adjusting the corresponding ratio formulas. Our model also allows temporal changes to viewing pressure of specific animal individuals/groups to be focused on by combining social media images with photo-identification records. Examples of this include analysis of specific dog dens (Cloutier *et al*., 2021), gorilla family groups (Molyneaux *et al*., 2021), and resident foraging sea turtles versus migratory turtles (Papafitsoros *et al*., 2021). This flexibility is particularly important because wildlife viewing pressure is not equally distributed across all animals present at a given site or time period. Certain individuals are often subjected to disproportionally higher viewing pressure, due to ecotourism activities incidentally or deliberately targeting these groups, particularly resident animals (Semeniuk *et al*., 2009; Christiansen and Lusseau, 2014; Schofield *et al*., 2015; Papafitsoros *et al*., 2021). Thus, social media could be used to tease out this information quantitatively, and introduce more appropriate watching practices and conservation measures.

The selected social media platform also influences interpretation (Ghermandi *et al*., 2020), particularly as the demographics of visitors and social media use change over time. Ideally, social media data should reflect actual human-animal interactions, while minimising user-induced biases, in parallel to revealing temporal variability in these interactions. For this reason, we used Instagram, because it captures real life human activity effectively (Tenkanen *et al*., 2017; Hausmann *et al*., 2018). This attribute allowed us to objectively identify temporal variation in viewing pressure. Alternatively, flickr has been widely used to infer spatial information on national park visitations, due to its easily accessible geotagged photographs (Wood *et al*., 2013; Barros *et al*., 2019; Ghermandi *et al*., 2020; Edwards *et al*., 2021); yet, its temporal correlation with ground-truthed data is lower than of Instagram (Tenkanen *et al*., 2017). Other platforms could also be used in our model, like YouTube (Otsuka and Yamakoshi, 2020; Taklis *et al*., 2020); however, larger time aggregation windows might be required to account for lower temporal correlation between the time of viewing and time of video uploading. In contrast, the demographics of Instagram users are not always representative of those of visitors to a focal site, i.e. generally younger users than other social media forms (Heikinheimo *et al*., 2017; Hausmann *et al*., 2018), and might also depend on country of origin (Ghermandi and Sinclair, 2019). Biases might also be self-diminishing, such as if, hypothetically, smartphone availability increased across years whereas Instagram (or any social media) use decreases. In this case, the net output of social media content might remain constant, even though the processes that govern information flow from the focal site to social media platforms are time dependent.

As with other technologies involving the remote collection of data, citizen science/social media data should be validated (or ground-truthed) using robust field data to guarantee conservation policies are informed appropriately (Jarić *et al*., 2020). Since information flow from the human-animal encounter to it being detectable in social media changes across sites, validation should be site-specific. The current study was methods-based, and so ground truthing was not the primary focus; however, our outputs closely aligned with previous studies at our site (Schofield *et al*., 2015; Papafitsoros *et al*., 2021). The ground-truthing of our model would involve estimating P, which would allow V_real_ to be directly estimated at a given time interval, rather than only the ratio in two time periods. This approach could provide a useful comparison of the ratio estimation we provide here; however, determining P remains challenging. P could be estimated by using questionnaires targeting visitors at focal sites, investigating whether visitors observed wildlife and uploaded any images on social media, or by quantifying actual viewing pressure using direct observations over time. Regular repetition of such surveys is necessary to determine the time dependence of P, particularly in the long term, i.e. across years.

## Conclusion

Most protected areas globally receive visitors that produce social media content related to them (Tenkanen *et al*., 2017), leading to the emergence of passive citizen science/iEcology/passive crowdsourcing. These records are typically used to study spatiotemporal visitation patterns and interaction of humans with the natural environment, and identify potential threats to wildlife (Sullivan *et al*., 2019; Jarić *et al*., 2020; Edwards *et al*., 2021; Papafitsoros *et al*., 2021; Molyneaux *et al*., 2021; Van Hamme *et al*., 2021; Cloutier *et al*., 2021). However, to accomplish this, social media-related biases and uncertainties must be overcome, with the current study making an important step towards accomplishing this. We modelled the flow of information from human-animal encounter to its appearance in social media, and inferred temporal changes in the number of encounters from temporal changes to corresponding social media imagery. We focused on quantifying uncertainty underlying such inferences, and identifying aggregation windows that combined increased temporal granularity to reduce uncertainty. We expect that continuous advances in automatic social media semantic data mining and machine learning will facilitate the creation of well-curated and meaningful datasets (Väisänen *et al*., 2021; Tuia *et al*., 2022), which combined with our framework will increase the number of studies using social media data to infer impacts on wildlife.

## Acknowledgments

We thank the Hellenic Civil Aviation Authority (CAA, http://www.ypa.gr) for providing passenger arrival data for the Zakynthos airport.

## Author contributions statement

Kostas Papafitsoros conceived the study and Lukáš Adam led the model development and the data analysis; Kostas Papafitsoros assimilated the social media data; Kostas Papafitsoros led the writing of the manuscript with critical contributions from Lukáš Adam and Gail Schofield.

## Supplementary material

### A. Derivation of the confidence interval (4)

This section derives the confidence interval (4). We recall that V_detected_ follows the binomial distribution with parameters V_real_ and P. Then its expectation is 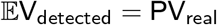. We start with the observation that the logarithm of the ratio of two independent binomial distributions rescaled by the number of trials, is approximately normal (Katz *et al*., 1978), provided the number of these trials is large, i.e.,

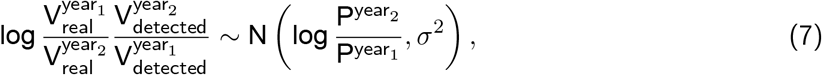

where

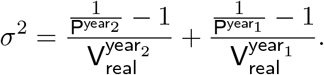

After reformulating *σ*^2^, we arrive at

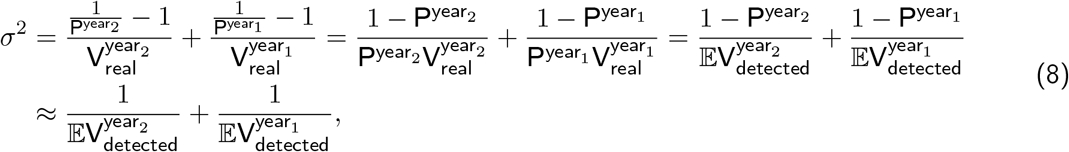

where the approximation in the second line of (8) assumes that the probabilities P^year_1_^ and P^year_2_^ are small. Since N(*μ*, *σ*^2^) = *μ* + *σ*N(0, 1), now (7) implies

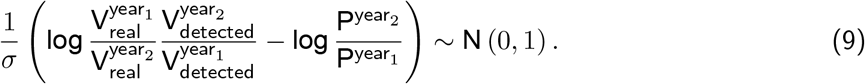

We fix an *α* ∈ (0, 1) describing the error in the confidence interval, which is typically *α* = 0.05, allowing for 5% error. We denote further Φ the cumulative distribution function of the standard normal distribution N(0, 1) and define 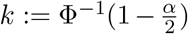. Thus, a sample from N(0, 1) lies in the interval [–*k, k*] with probability 1 – *α*. Applying this result to (9) means that both inequalities in

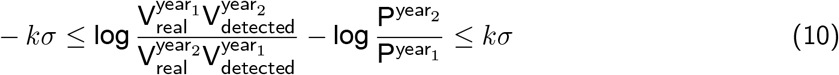

are satisfied with probability 1 – *α*. Formula (10) can now be reformulated as

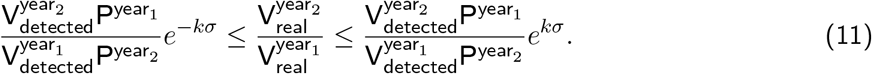

By combining (11) with (8) and replacing the expectations by the realisation of the random variables, we derive the confidence interval (4).

### B. Uncertainty quantification and additional assumptions

To derive the confidence interval (4), we made three additional assumptions:

i. The number of the real viewing pressures 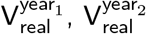 is large. This is required for the approximation of the distribution of the ratio of two binomial distributions as in (7).
ii. The probabilities P^year_1_^, P^year_2_^ are small. This is required to replace both 1 – P^year_1_^ and 1 – P^year_2_^ by 1 in (8).
iii. The observed values 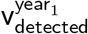 and 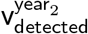 are close to the respective means 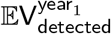 and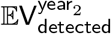. This is required when using (8) to derive (4) in the last step above.

Assumption (i) is satisfied for large enough time duration, and/or when multiple animal viewings are considered. Assumption (ii) is expected to be satisfied, if the number of real observations is much higher that the corresponding number of detected entries. Assumption (iii) is typically not satisfied exactly due to the nonzero variance of 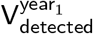 and 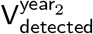, i.e., due to their randomness. However, validation experiments using simulations indicate that the introduced error from this approximation is small.

### C. Multiplicative bounds in Figure 8

One can easily observe the symmetry of the confidence interval (4) derived above, that is upon switching the two years year_1_ and year_2_, we obtain

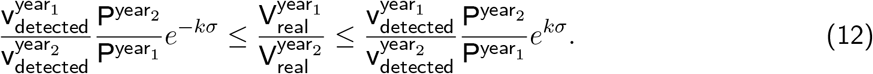

If we computed the standard (additive) width of the confidence interval, the results would change for (4) and (12). However, by computing the “multiplicative” bounds, i.e. dividing the upper bound by the point estimate, we obtain *e^kσ^* for (4) and (12). This is desirable, as switching the order of years should not change results. This is the reason for using multiplicative instead of additive bounds in Figure 8.

### D. Validation of the derived confidence intervals

When we derived the confidence intervals, we used several simplifying assumptions. We commented on them in Supplementary material B and provided their numerical justification in Figure 2. We describe now the exact process of generating this figure. We selected a series of combination of detected values 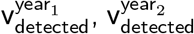, fixed the probabilities P^year_1_^ = P^year_2_^ and computed the confidence interval according to (4). For this computation we only need to know the ratio P^year_2_^/P^year_1_^ and not the individual probabilities.

To verify the confidence intervals, we reverse the generating process. We recall that V_detected_ follows the binomial distribution with the number of samples V_real_ and selection probability P. We therefore sample the individual samples with probability P until vdetected successes is reached. This process is equivalent to sampling from the negative binomial distribution. In this way, we generate V_real_ and then compute the ratio 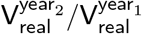. We repeat this process 10,000,000 times, ignoring the fraction of 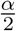 observations from both sides. The range of the remaining trials gives us the real confidence interval on level *α*. Having our and the true confidence interval, we compute their IoU metric, which is their intersection divided their union. This is depicted in Figure 2.

### E. Supplementary tables

**Supplementary table 1.** Numbers of boat entries for May-October for 2019-2021.

|  | May | Jun | Jul | Aug | Sep | Oct | Total |
| --- | --- | --- | --- | --- | --- | --- | --- |
| 2019 | 53 | 257 | 360 | 473 | 211 | 28 | 1382 |
| 2020 | 4 | 5 | 83 | 196 | 78 | 21 | 387 |
| 2021 | 14 | 98 | 241 | 356 | 131 | 37 | 877 |

**Supplementary table 2.** Monthly numbers of airport arrivals to Zakynthos for May-October for 2019-2021, according to the Hellenic Civil Aviation Authority (CAA) (http://www.ypa.gr).

|  | May | Jun | Jul | Aug | Sep | Oct | Total |
| --- | --- | --- | --- | --- | --- | --- | --- |
| 2019 arrivals | 87938 | 168066 | 212384 | 208821 | 152769 | 29976 | 859954 |
| 2020 arrivals | 249 | 988 | 58561 | 102481 | 39240 | 6994 | 208513 |
| 2021 arrivals | 8506 | 52172 | 136629 | 158328 | 107323 | 30648 | 493606 |

**Supplementary table 3.** Global number of Instagram users. Source: https://www.statista.com/statistics/183585/instagram-number-of-global-users, (*accessed on 14.01.2022*)

|  | 2019 | 2020 | 2021 |
| --- | --- | --- | --- |
| Number of users (millions) | 814.5 | 1000.8 | 1074.3 |

## Footnotes

1 https://www.statista.com/statistics/183585/instagram-number-of-global-users, (*accessed on 14.01.2022*)

2 https://uk.mathworks.com/matlabcentral/fileexchange/99404-conservative-regridding

